# Cytogenetic Diversity and Genome Size Variation in *Cuscuta* L. subgenus *Grammica* section *Subulatae* (Convolvulaceae): Insights from *C. argentinana* Yunk. and *C. parodiana* Yunk

**DOI:** 10.1101/2025.08.21.671489

**Authors:** Amalia Ibiapino, Andrea Pedrosa-Harand, Juan Urdampilleta

## Abstract

The genus *Cuscuta* exhibits remarkable cytogenetic diversity, strongly influenced by heterochromatin dynamics. The genus is divided into four subgenera, the subgenus *Grammica* is almost exclusively found in the Americas, and South America is the first major diversification center. The section *Subulatae* consists of species primarily found in Argentina. To cytogenetically characterize the *Cuscuta* species of *Subulatae* and investigate the chromosomal evolution of the subgenus *Grammica*, CMA/DAPI banding, FISH with 5S and 35S rDNA, and flow cytometry were performed on *C. argentinana* and *C. parodiana*. Both species exhibited 2*n* = 30 with differences in chromosome size, heterochromatic banding patterns, and genome size. *Cuscuta argentinana* has smaller chromosomes and genome (1C = 1.49 Gbp) and a higher number of metacentric chromosomes and CMA⁺/DAPI⁻ bands, whereas *C. parodiana* (1C = 2.79 Gbp) exhibits heterochromatin accumulation and a higher number of submetacentric chromosomes and heterochromatin accumulation. While the number of 5S rDNA sites was the same (six sites), *C. parodiana* presented one more pair of 35S rDNA sites. The results suggest that genome variation in *Cuscuta* section *Subulatae* is associated with heterochromatin amplification. These findings contribute to understanding of *Grammica* diversification and the role of heterochomatin in the chromosomal evolution of the genus.

## Introduction

The genus *Cuscuta* L. (Convolvulaceae) comprises parasitic plants that exhibit great cytogenetic diversity. With a basic chromosome number of *x* = 15, the genus stands out for its species with highly variable chromosome numbers (ranging from 2*n* = 8 to 2*n* = 150) (Ibiapino et al. 2022a), karyotypes ranging from symmetrical to bimodal (Ibiapino et al. 2022a; Ibiapino et al. 2022b), and both holocentric and monocentric chromosomes (García and Castroviejo 2002; Guerra and García 2004). Additionally, *Cuscuta* displays a remarkable 128-fold genome size variation, from the smallest genome (1C = 0.27 Gbp in *C. australis* R. Br.) to the largest (1C = 34.73 Gbp in *C. reflexa* Roxb.) (García et al. 2014; Neumann et al. 2021; Ibiapino et al. 2022a). Beyond the role of polyploidy, the dynamics of heterochromatin organization in these species appear to strongly influence the karyotypic variation observed in the genus. Giant genomes, such as that of *C. monogyna* Vahl (1C = 32.45 pg) (Ibiapino et al. 2020), show significant amplification of various classes of repetitive DNA (Neumann et al. 2021). Furthermore, the accumulation of repetitive DNA has been identified as one of the main factors responsible for the emergence of bimodal karyotypes in the subgenus *Pachystigma* (Engelm.) Baker & C. H. (Ibiapino et al. 2022).

The genus *Cuscuta* is highly cosmopolitan, with each of its four subgenera occupying distinct geographic regions. The subgenus *Monogynella* (Des Moul.) Peter, Engl. & Prantl is found in southern and eastern Asia, Europe, and Africa, with one species, *C. exaltata* Engelm., native to southeastern North America. The subgenus *Cuscuta* is native to Europe, Africa, and Asia, although some species have been introduced and naturalized in the Americas, Australia, and New Zealand. The subgenus *Pachystigma* consists of only five species, all endemic to South Africa and is sister to *Grammica* (Lour.) Yunck. *Grammica* is distributed almost exclusively in the Americas (García et al. 2014; Costea et al. 2015). This last subgenus harbours species with greater variation in genome size, chromosome number, and chromosome morphology, as well as cases of auto- and allopolyploidy (García et al. 2018; Ibiapino et al. 2022a; Ibiapino et al. 2025). Additionally, within *Grammica*, the *Subulatae* section (clade O) is one of the largest clades and represents the first major diversification event within the subgenus. This section includes species occurring in South America, and as *C. argentinana* Yunk. and *C. parodiana* Yunk. occuring in Argentina (García et al. 2014; Costea et al. 2015). Species of the genus *Cuscuta* parasitize at least 25 different crops and pose a problem in at least 55 countries (Kaiser et al. 2015). Despite the losses, the genus *Cuscuta* serves as a model for studies on haustorium development and chromosomal evolution, as it includes species with key cytogenetic characteristics of interest (Jhu and Sinha 2022; Ibiapino et al. 2022).

Until now, published cytogenetic data for clade O species are limited to chromosome counts, for six species: *C. purpurata* Phil., *C. chilensis* Ker Gawl. and *C. grandiflora* Kunth (2*n* = 30) *C. parodiana* (2n = 30 and 60) (Paéz et al. 2011; Andrada et al. 2018; García et al. 2019), *C. globiflora* (n = 48) and *C. cristata* (2n = 14) (Páez et al. 2011) and genome sizes, for only two species, *C. purpurata* (1C = 2.96 Gbp) and *C. chilensis* (1C = 2.8 Gbp) (McNeal et al. 2007; Ibiapino et al. 2022). This represents a low sample size, considering that the *Subutatae* section includes 31 species (Costea et al. 2015). More in-depth karyotypic characterizations will help to understand the evolutionary mechanisms from the base of *Grammica* to the diversification of the other species. The subgenus *Grammica* is the largest and most geographically widespread subgenus (García et al. 2014). The extensive cytogenetic variation found in the genus may be linked to ecological factors, particularly its parasitic lifestyle, which allows these plants to escape certain environmental constraints. This lifestyle facilitates the colonization of large areas and promotes specific genomic changes (Gruner et al. 2010; Piednoël et al. 2012).

One of the key strategies for karyotypic characterization is the use of CMA/DAPI banding. This technique employs two fluorochromes with different affinities to identify the number and position of heterochromatic GC-rich bands (Chromomycin A3 – CMA) and AT-rich bands (4’,6-diamidino-2-phenylindole – DAPI) (Cabral et al. 2006). Heterochromatin is mainly composed of repetitive DNA sequences, which can be organized in *tandem* or dispersed throughout the genome (Heslop-Harrison and Schwarzacher 2011). Among the *tandem* sequences, satellite DNA and ribosomal DNA (5S and 35S) stand out, with the rDNA being highly conserved and useful for phylogenetic comparisons between different species (Roa and Guerra 2012, 2015). The diversity and abundance of these sequences may be linked to environmental interactions, potentially explaining why species occupying temperate and tropical areas exhibit genome size differentiation among closely related taxa (Biscotti et al. 2019; Van-Lume et al. 2017, 2019). As previously mentioned, the organization of the repetitive fraction of the genome in many *Cuscuta* species is closely associated with the genus’s karyotypic diversity. Considering the subgenus *Grammica* distributed and the diversification of the Clade O, these data suggest that genomic changes that occurred after the diversification event in clade O may have contributed to the genomic diversity found in the subgenus *Grammica*.

Therefore, this study aims to provide more detailed cytogenetic data on *C. argentinana* and *C. parodiana*, two species from clade O in South America, using CMA/DAPI banding, fluorescent in situ hybridization (FISH) with 5S and 35S rDNA probes, and genome size estimation via flow cytometry. This study seeks to address the following questions: (1) How do heterochromatin distribution patterns in *C. argentinana* and *C. parodiana* relate to their karyotypic evolution and genome size variation? (2) Do *C. argentinana* and *C. parodiana* share karyotypic characteristics that support the definition of the section *Subulatae*?

## Material and methods

### Material

The species studied were collected in the Catamarca and Tucuman provinces of Argentina and vouchers were included in the Botanical Museum of Córdoba herbarium (CORD) (Table 1).

**Table 1:** Collection data, chromosome number, karyotypic formula, average size of the largest and smallest pair, total haploid length and genome size (1C) of *Cuscuta argentinana* and *Cuscuta parodiana*.

| Specie | 2n | karyotypic formula | largest and smallest pair size (μm) | total haploid length (μm) | genome size (Gbp) | Voucher |
| --- | --- | --- | --- | --- | --- | --- |
| *Cuscuta argentinana* | 30 | 28M + 2SM | 5.51 - 1.92 | 79.96 | 1.49 | ARG, Catamarca, *Urdampilleta 914 (CORD)* |
| *Cuscuta parodiana* | 30 | 6M + 24SM | 4.14 - 2.18 | 95.92 | 2.79 | ARG, Tucuman, *Urdampilleta 917 (CORD)* |

### Slides preparation, sequential chromosome banding and FISH, and chromosome length measurement

Slide preparation was performed using apical meristems from adult plants cultivated in the experimental garden of the IMBIV, growing in *Baccharis* sp. as host. The material was pretreated with 8-hydroxyquinoline for 6 h at 18 °C for *C. argentinana* and 24 h at 10 °C for *C. parodiana*, fixed in 3: 1 (v/v) ethanol: acetic acid for 2-24 h at room temperature, and stored at -20 °C. Subsequently, the material was washed in distilled water, digested in Pectinex (Novozimes), and squashed in 60% acetic acid. Double CMA/DAPI staining was performed as described in Ibiapino et al. (2022a). The images were captured with an Olympus BX61 microscope coupled with a monochromatic camera and Cytovision software (Leica Biosystems). The images were pseudo-coloured using Adobe Photoshop CS3 software version 10.0. After image capture, slides were destained for 30 min in Carnoy and 1 h in absolute ethanol and stored at -20 °C.

The destained slides were subjected to FISH according to the protocol detailed by Pedrosa et al. (2002), with the following modifications, the slides containing the hybridization mix were placed in the thermocycler for the following cycle: 90°C for 10 min, 48°C for 10 min, and 38°C for 5 min. They were then incubated in a preheated humidity chamber at 37°C overnight. After incubation, post-hybridization washes were performed under agitation in the following order: a 2×SSC wash at room temperature until the coverslip detached, a 2×SSC wash at 42°C for 10 min, a 0.1×SSC wash at 42°C, a 2×SSC wash at 42°C for 10 min, a 4×SSC/0.2% Tween wash at 42°C for 10 min, and finally, a 4×SSC/0.2% Tween wash at room temperature. Subsequently, the slides were stained with 10 µl of DAPI in a humid chamber at room temperature for 20 min, washed with 4×SSC/0.2% Tween, and mounted with 10 µl of DABCO.

Two rDNA probes were used: for the 5S rDNA, the four PLOPs end-labelled with Tamra, and for the 35S rDNA, the 12 PLOPs end-labeled with FAM (Macrogen), as described by (Waminal et al. 2018). The FISH pictures were obtained as previously described.

Chromosome size estimations were based on measurements of the ten best metaphases of each species, using Adobe Photoshop CS3 software. Chromosome arm ratio was used to classify chromosomes as metacentric or submetacentric according to Ibiapino et al. (2020). To assemble the karyogram, the chromosomes were organized by size, from largest to smallest, also considering the presence of CMA/DAPI bands and rDNA sites for identifying homologous chromosomes.

### Flow cytometry

A suspension of nuclei from shoot tips cultivated in the experimental garden of the Federal University of Pernambuco was prepared using WPB buffer (Loureiro et al. 2007). The nuclei were stained using propidium iodide and the amount of nuclear DNA was estimated using the CyFlow SL flow cytometer software (Partec, Görlitz, Germany). *Solanum lycopersicum* L. “Stupické polní rané” (1C = 0.96 pg) was used as internal standard (Doležel 1991). The final genome size was based on three different measurements with 5,000 nuclei each, and using the equation “(Sample peak mean/Standard peak) × mean 2C DNA content of internal control (pg)” and the software FloMax (Partec) for data processing. To present these results the genome sizes were converted to Gbp.

## Results

### Chromosome number, size and morphology, and DNA amount

Both *C. argentinana* and *C. parodiana* exhibited 2*n* = 30 (Figure 1), with some differences in chromosome size and morphology, heterochromatic band distribution pattern, and genome size. In *C. argentinana*, chromosome size varied 2.78-fold between the largest (5.31 μm) and the smallest (1.91 μm) chromosome pairs. The total haploid length of this species was 79.96 μm (Figure 2, Table 1), and its genome size was 1C = 1.49 Gbp (Table 1). Only the largest chromosome pair of *C. argentinana* exhibited a submetacentric morphology, while all others were metacentric (Figure 2) presenting a karyotypic formula 28M + 2SM.

**Figure 1:**
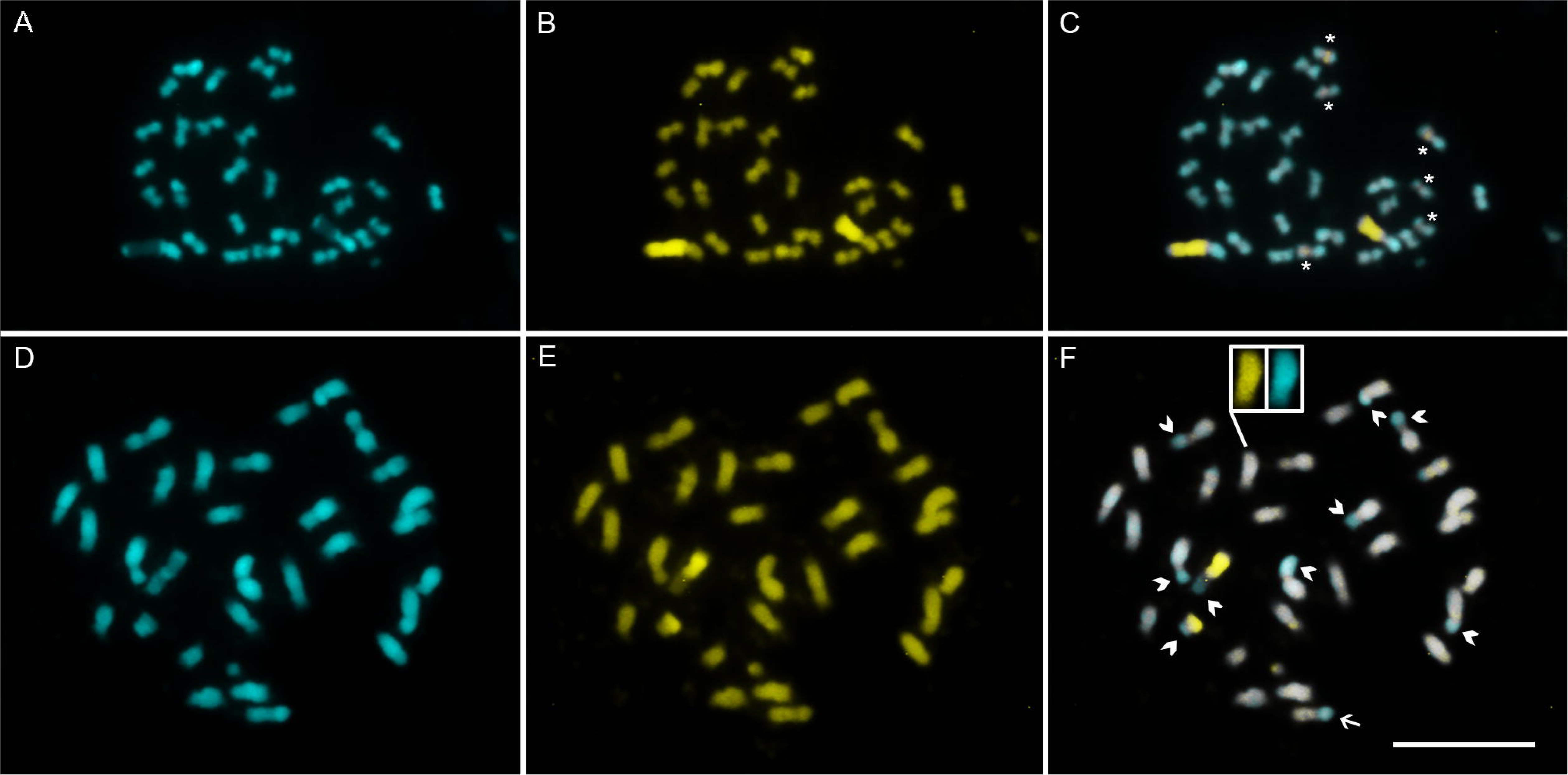
Mitotic metaphases of *Cuscuta argentinana* (A, B and C) and *Cuscuta parodiana* (D, E and F) showing the results of DAPI (blue) in A and D; CMA (yellow) in B and E; and merge pictures in C and F. The asterisks in C show the smaller CMA⁺/DAPI⁻ bands of *C. argentinana* and the arrows head in F show the CMA⁰/DAPI⁺ bands on the short arms of some *C. parodiana* chromosomes. Inset in F showing a chromosome that has no evident peri/centromeric gap either with CMA staining or with DAPI staining. Bar in G representing 10μm.

**Figure 2:**
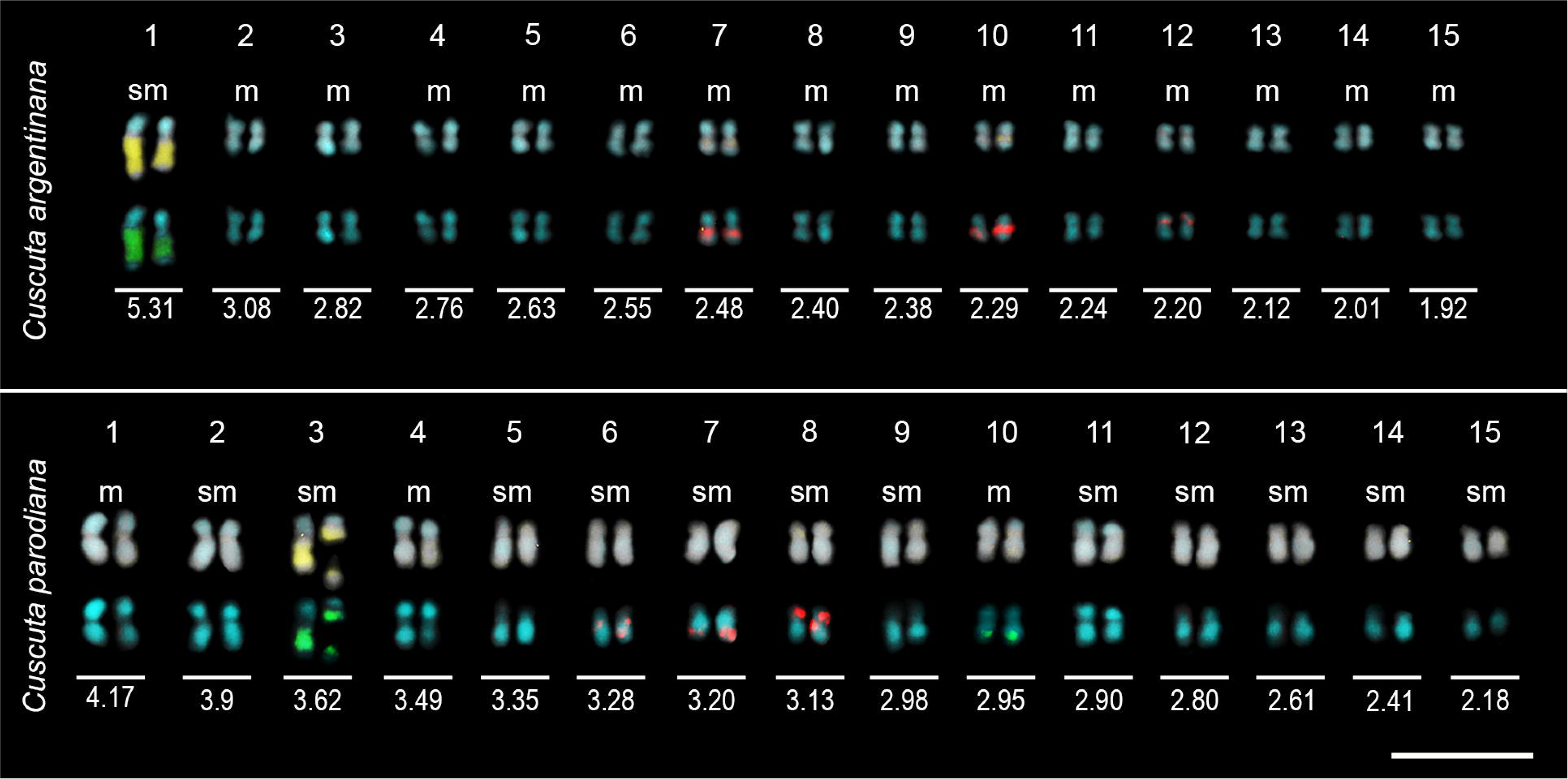
Karyograms of *Cuscuta argentinana* and *Cuscuta parodiana* showing the morphology and average size of each chromosome pair. The average size is based on all cells measured, the chromosomes shown in the karyogram were taken from the metaphases shown in Figure 1. Chromosomes are arranged in descending order of size, from largest to smallest. The top rows display chromosomes stained with CMA (yellow) and DAPI (blue). The bottom rows show chromosomes subjected to FISH with 5S (red) and 35S (green) rDNA probes. Scale bars in the lower right corners represent 10 μm.

In *C. parodiana*, chromosome size varied 1.91-fold between the largest pair (4.17 μm) and the smallest pair (2.18 μm). Its total haploid length was 95.92 μm (Figure 2, Table 1), and its genome size was 1C = 2.79 Gbp (Table 1). In this species, most chromosome pairs (a total of 12) exhibited a submetacentric morphology, while three pairs (one, four, and ten) were metacentric; its karyotypic formula is 6M + 24SM.

### CMA/DAPI bands and rDNA sites

*Cuscuta argentinana* exhibited eight CMA⁺/DAPI⁻ bands (Figure 1 A-C), with a larger band in the pair 1 forming the nucleolus organizing region (NOR) or secondary constriction. Smaller bands were present in the pairs 7, 10 and 12 (Figure 2 C, Figure 3). All chromosomes displayed a neutral peri/centromeric region for both CMA and DAPI, resulting in a very distinct gap (Figure 1 A, B and C). Regarding the rDNA sites, this species had three pairs of 5S rDNA sites (Figure 3 A) colocalized with the smaller CMA⁺/DAPI⁻ bands (Figure 2 and 3). Additionally, a pair of 35S rDNA sites was present on the long arm of the largest chromosome pair, colocalized with NOR (Figure 2 C and 3 A).

**Figure 3:**
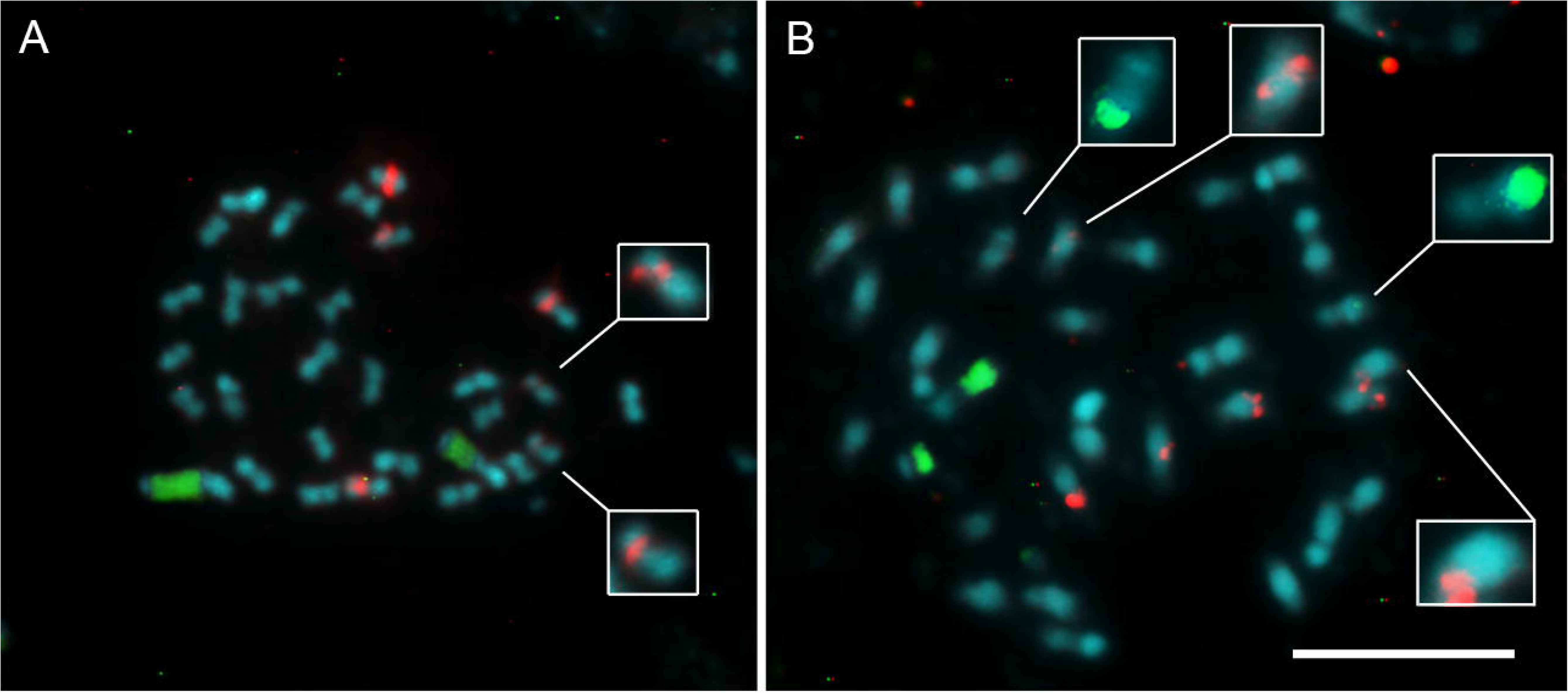
Mitotic metaphases of *Cuscuta argentina* (A) and *Cuscuta parodiana* (B) with FISH using 5S (red) and 35S (green) rDNA. InseRts show fainter rDNA sites with increased scale and exposure to allow visualization. Bar in B represents 10μm.

*Cuscuta parodiana* exhibited CMA⁰/DAPI⁺ bands on the short arms of 10 chromosomes (Figure 1 D, E and F), corresponding to pairs 1, 2, 3, 4, and 11 (Figure 2 F). Only one CMA⁺/DAPI⁻ band was identified, located on the long arm of pair three, forming a secondary constriction. Overall, the chromosomes of this species were strongly stained for both CMA and DAPI (Figure 1 F). The peri/centromeric region of this species was not as evident in all chromosomes, sometimes stained with both CMA and DAPI, as observed in pair 7 (Figure 2). This species also had six 5S rDNA sites: on the long arms of chromosome pairs 6 and 7 and on the short arm of pair 8 (Figure 3 B). Regarding the 35S rDNA sites, *C. parodiana* presented two pairs, one on the long arm of chromosome pair 3 and another on the long arm of pair 10. Only the largest 35S rDNA site is colocalized with CMA⁺/DAPI⁻ band in pair 3 (Figure 2 F, Figure 23.

## Discussion

### Chromosome number variation

Considering that the basic chromosome number of *Cuscuta* is *x* = 15 (Ibiapino et al., 2022a), the accessions of *C. argentinana* and *C. parodiana* analysed in the present study are diploid, both with 2*n* = 30. Previous studies have reported intraspecific chromosome number variation in *C. parodiana*, with accessions containing *n* = 15 and *n* = 30 (Andrada et al. 2018). Intraspecific numerical variation has also been reported for other species of *Cuscuta*. In the holocentric species, *C. epithymum* L. (subgenus *Cuscuta*), for example, *n* = 7, 8, 14, 15, and 16 have been documented (García and Castroviejo 2002; Marhold et al. 2019). In the subgenus *Grammica*, for *C. denticulata* Engelm., the reported chromosome numbers are *n* = 15 and 30, with variation occurring due to autopolyploidy (García et al. 2018). This phenomenon can result from several factors, one of which is the fusion of unreduced gametes, caused by errors during meiosis (Goulet et al. 2017). Analyses carried out by Andrada et al. (2018) revealed some irregularities in the meiosis of certain accessions of *C. parodiana*, such as the formation of trivalents and tetravalents, chromatin bridges, chromosomes outside the metaphase plate, and cytomixis. This could explain the numerical variation found in the karyotype of *C. parodiana*.

*Cuscuta purpurata*, *C. chilensis*, and *C. grandiflora*, which belong to the same section, also exhibit 2*n* = 30 (García et al. 2019). However, considering that this section is the largest within the subgenus *Grammica*, comprising 31 species (Costea et al. 2015), the current sample size remains too small to conclude that the chromosome number in this clade is conserved. Other representatives, such as *C. cristata* (*n* = 14) and *C. globiflora* (*n* = 48) (Paéz et al. 2011), have less commonly reported chromosome numbers in *Cuscuta*, indicate that mechanisms responsible for numerical chromosome variation—such as autopolyploidy and allopolyploidy, which are observed in other clades of *Grammica*, or descending dysploidy, which is observed mainly in the holocentrics from subgenus *Cuscuta* (García et al. 2018; Ibiapino et al. 2022a; Ibiapino et al. 2025) —may also be present in the *Subulatae* section.

### Genome size variation in *Subulatae* section

Despite belonging to the same section (García et al. 2014), *C. argentinana* and *C. parodiana* exhibit significant differences in chromosome morphology and genome size. *Cuscuta parodiana* has larger chromosomes, with more submetacentric pairs than *C. argentinana*. The total haploid length varies 1.2-fold between *C. argentinana* and *C. parodiana*, corresponding to the difference in genome size between these species of 1.87-fold. Other known cases of genome size in this section are 1C = 2.96 Gbp in *C. purpurata* and 1C = 2.8 Gbp (Ibiapino et al. 2022a) in *C. chilensis* (McNeal et al. 2007). Both species are diploid and have genome sizes similar to *C. parodiana*. Considering that the average genome size in the clade O species is 1C = 2.51 Gbp and *Cuscuta parodiana*, *C. chilensis*, and *C. purpurata* exhibit genome sizes similar to each other, there is a reduction in genome size in *C. argentinana* compared to other species of this clade. These differences may have arisen due to genomic changes following the diversification of the subgenus *Grammica*. These species belong to the *Subulatae* section (Clade O), the first major diversification clade of the subgenus (García et al. 2014; Costeae et al. 2015). This subgenus extends throughout the Americas and occupies diverse ecological niches (García et al. 2014).

It is known that variations in the geographic distribution and ecological niche patterns can greatly influence the distribution of heterochromatin in plants (Van-Lume et al. 2017, 2019). Ecological factors can lead, for example, to the expansion or contraction of various families of repetitive DNA that constitute this heterochromatin, which in turn can generate genome size variability within related groups of organisms (López-Flores and Garrido-Ramos 2012; Bourque et al. 2018). Although *C. chilensis* and *C. purpurata* occur mainly in Chile and Peru, respectively, and *C. argentinana* and *C. parodiana*, analysed in this study, are geographically separated by the mountain range of Aconquija National Park, the variation in genome size in *Cuscuta* appears to be associated with its parasitic lifestyle.

An ancestral character reconstruction in *Cuscuta* suggested an ancestral genome size of ca. 1C = 12 Gbp (Ibiapino et al. 2022a), while the genome size average in other Convolvulaceae is approximately 1C = 0.97 (Ibiapino et al. 2022a). When analyzing only the subgenus *Grammica*, some of its sections exhibit a tendency toward genome size expansion, while others undergo significant reduction. In North America, for example, the diversification of this subgenus followed opposite trends: while the sections *Oxycarpae* (clade D) and *Denticulatae* (clade E) show an increase in genome size, other clades, such as *Californicae* (clade A) and *Cleitogrammica* (clade B), have the smallest genome sizes within the genus (Ibiapino et al. 2022a).

There is a tendency for parasitic plants to develop larger and more complex genomes. In *Cuscuta* some species can be classified as holoparasites, while another species can be classified as hemiparasite according to the degrees os plastome reduction (Garcia et al. 2014; Banerjee and Stefanović 2020) The plastid genome tends to be conserved in autotrophic plants but undergoes reduction in parasitic plants. The fact that *Cuscuta* exhibits varying degrees of plastome reduction and loss of photosynthesis-related genes results in some species of the genus being classified differently (Banerjee and Stefanović 2019, 2020). The increase of genome size in *Cuscuta*, compared with other Convolvulaceae, can be attributed to parasitic lifestyle, which can eliminate constraints associated with DNA accumulation and replication because the restrictions imposed by the growth rate of meristem or the “genomic economy”, as they obtain resources from their hosts (Gruner et al. 2010; Piednoël et al. 2012). Species of section *Subulatae* are holoparasitic (Braukmann et al., 2013), compared to the ancestral genome size of the genus *Cuscuta*, the clade O shows a reduction in genome size, but considering the subgenus *Grammica*, this clade present genome sizes that are more intermediate between the smallest (*C. australis* – 1C = 0.27 Gbp) and the largest (*C. indecora* - 26.39) within the subgenus (Ibiapino et al. 2022). In Orobanchaceae, which has another holoparasitic plant genera, such as *Orobanche* L. and *Phelipanche* Pomel have the largest genomes compared to hemiparasitic and autotrophic genera within the same family (Gruner et al. 2010; Piednoel et al. 2012). The different degree of plastome reduction in *Cuscuta* can explain the variation in the genome size found in the genus, suggesting that this variation is more closely linked to ecological factors, such as parasitism, than to the geographic distribution of the species.

### Heterochromatin distribution and rDNA sites

*Cuscuta argentinana* presents a larger number of CMA⁺/DAPI⁻ bands; however, *C. parodiana* appears to exhibit a strong accumulation of repetitive DNA in the long arms of the chromosomes, which are equally stained by both CMA and DAPI. These data suggest that the difference in genome size is influenced by variations in heterochromatin patterns, which, in turn, may differ due to the tendency of parasitic plants to develop more complex genomes. This heterochromatin accumulation in the long arms of the *C. parodiana* chromosomes may have led to the elongation of these arms, resulting in the predominance of a submetacentric chromosome morphology in this species. Additionally, ten of these chromosome pairs exhibit an accumulation of DAPI⁺ regions, particularly in the short arms. Further evidence of heterochromatin accumulation in *C. parodiana* includes the peri/centromeric region, which appears more intensely stained, and the presence of an additional pair of 35S rDNA sites compared to *C. argentinana*. In *C. argentinana*, the peri/centromeric region appears as a distinct gap, and this species possesses only a single pair of 35S rDNA sites. The number and position of rDNA sites in *Cuscuta* can vary considerably. Some species, such as *C. howelliana* P. Rubtzov (subgenus *Grammica*), have a single pair of 5S rDNA sites and a single pair of 35S sites, whereas *C. monogyna* (subgenus *Monogynella*) shows the highest known numbers, with 18 pairs of 5S rDNA sites and 15 pairs of 35S sites (Ibiapino et al. 2022).

The subgenus *Pachystigma*, is restricted to South Africa and sister to *Grammica*, consists of five species (García et al. 2014), but rDNA sites are only known for *C. nitida* E.Mey. ex Choisy. This species has a single pair of 5S rDNA sites, which occupy almost the entire short arm of its largest chromosome pair, along with two 35S rDNA sites in nearby regions (Ibiapino et al. 2022b). Additionally, one of the 35S pairs is found on the same chromosome as the 5S rDNA site (Ibiapino et al. 2022b). *Cuscuta argentinana* and *C. parodiana* appear to have undergone an expansion in the number of 5S rDNA sites compared to *Pachystigma*, as both species have three pairs. Regarding 35S rDNA sites, these two species seem to have undergone significant rearrangements compared to *C. nitida*, since none of their 35S sites are adjacent to 5S sites, and in *C. parodiana*, one of the rDNA 35S pair is located in a more terminal region.

Many species exhibit adjacent rDNA sites, meaning they occur on the same chromosome, as seen in *C. denticulata*, *C. partita* Choisy, and the previously mentioned *C. nitida*, *C. howelliana* and *C. monogyna* (Ibiapino et al. 2019, 2020, 2022a). In *C. globosa*, which belongs to the *Gracillimae* section (clade N) and diversified later than the *Subulatae* section, the number and position of sites are similar to those reported in *C. parodiana* (García et al. 2014; Ibiapino et al. 2022a), suggesting that the absence of adjacent rDNA sites may be a derived condition which arises independently in certain *Cuscuta* clades. However, in the following clade, the *Indecorae* section (clade M), which is sister to *Gracillimae* (clade N), *C. indecora* presents five pairs of 5S rDNA sites and two pairs of 35S rDNA sites, indicating an expansion of the 5S sites (Ibiapino et al. 2020). The presence of one pair of adjacent sites suggests an independent rearrangement associated with the expansion of rDNA sites (Ibiapino et al. 2020), contributing to the increase in genome size (Neumann et al. 2021).

It is well established that heterochromatin can accumulate in pericentromeric regions, as well as in terminal and interstitial chromosomal domains, generating differences even among closely related species (Biscotti et al. 2015). Kao (2001) characterized nine species of the orchid genus *Phalaenopsis* Blume and found that both genome size and heterochromatin banding patterns varied. The authors demonstrated that the differential accumulation of AT-rich heterochromatin was the primary driver of karyotypic variation in orchids and suggested that the amplification of these sequences could contribute to speciation. This study suggests that *C. parodiana* has a greater accumulation of heterochromatin than *C. argentinana*, which may have contributed to the difference in genome size reported here.

### Heterochromatin and karyotype symmetry

With the increasing availability of sequenced plant genomes, it has been possible to observe a substantial accumulation of repetitive DNA over time in these organisms. Such changes can drive chromosomal structural rearrangements and influence plant karyotype evolution (Vimala et al. 2021). In other *Cuscuta* species (except the subgenus *Pachystigma*) where heterochromatin expansion has been reported, the karyotypes remain symmetrical (García et al. 2019, Ibiapino et al. 2020; Neumann et al. 2021). Until now there is no data available on the genome size of any species within subgenus *Pachystigma*. However, the total haploid chromosome length reported for *C. nitida* is 107.6 µm (Ibiapino et al. 2022b) suggests a larger genome size compared to *C. argentinana* (79.96 µm) and *C. parodiana* (95.92 µm). The subgenus *Pachystigma* includes species characterized by bimodal karyotypes, a trait that arose due to the differential accumulation of distinct classes of repetitive DNA in only two chromosome pairs (García et al. 2019; Ibiapino et al. 2022b). This accumulation led to an increase in the size of these pairs relative to the others, leading to genome size variation between these two sister species.

The maintenance of karyotypic symmetry despite the amplification of repetitive sequences has already been reported in another *Grammica* species, *C. indecora*. This species exhibits intraspecific genome size variation (2C = 45.58, 50.03, and 65.54 pg), which is significantly larger than the genome sizes reported for other *Grammica* species (around 1C = 2.45 Gbp) (Ibiapino et al. 2020; Ibiapino et al. 2022a). The expansion of its genome size was driven by the amplification of repetitive sequences. The repetitive fraction of this species constitutes 43.29% of its genome (Neumann et al. 2021); however, its karyotypic symmetry was preserved because this amplification occurred relatively evenly across all chromosomes. *C. indecora* exhibits both CMA⁺/DAPI⁻ and CMA⁻/DAPI⁺ bands distributed across all chromosomes, mainly in interstitial and pericentromeric regions (Ibiapino et al. 2020). A similar process may be occurring in clade O with *C. argentinana* and *C. parodiana*. The accumulation of heterochromatin appears to be uniform across the entire chromosome set, which has contributed to the maintenance of karyotypic symmetry in clade O. Further analyses involving next-generation sequencing, characterization of the repetitive fraction, and mapping of the most abundant sequences on chromosomes via FISH will be crucial to understanding how heterochromatin amplification contributes to the karyotypic symmetry of *C. argentinana* and *C. parodiana*.

The largest chromosome pair of *C. argentinana* is noticeably larger than the others, but this does not classify the species as bimodal. The difference between the largest pair (5.35 µm) and the average size of the smaller chromosome set (1.92 µm) is 2.78-fold. Nonetheless, this species exhibits a gradual decrease in chromosome sizes, with the difference between the first and second pairs being only 1.72-fold. In a truly bimodal karyotype, the largest chromosome pair must be at least twice the size of the smallest set (McKain et al. 2012). Bimodality resulting from the uneven accumulation of repetitive sequences, like occur in the subgenus *Pachystigma*, has also been documented in other genera. In *Eleutherine bulbosa* (Mill.) Urb., for instance, bimodality emerged due to the differential accumulation of various classes of repetitive DNA, which, although present in all chromosomes, were significantly enriched in the largest chromosome pair (Báez et al. 2019). In *Cuscuta*, the transition to bimodal karyotypes is a condition that characterizes the subgenus *Pachystigma*. The karyotype of *C. argentinana* is symmetrical, as is that of *C. parodiana*, where the chromosome pairs are more similar in size and exhibit a gradual decrease from the largest to the smallest pair.

This study revealed significant differences in chromosome organization and genome size between *C. argentinana* and *C. parodiana*, suggesting that heterochromatin distribution played a crucial role in the karyotypic differentiation of these species. Despite the variation in genomic content and CMA/DAPI banding patterns, both species maintain karyotypic symmetry, indicating that the expansion of repetitive sequences occurred in a relatively uniform manner. This pattern is consistent with other species of the genus *Cuscuta*, except for the subgenus *Pachystigma*. These data reinforce that the parasitic lifestyle of the genus contributes to complex genome organization, leading to variations in genome size even among species of the same clade. *Cuscuta* and highlight the need for additional studies, including the analysis of repetitive sequences through next-generation sequencing (NGS) techniques. The integration of these data will provide a more comprehensive view of the genomic mechanisms involved in the diversification of the subgenus *Grammica* and will contribute to the understanding of the relationship between genome size evolution and the parasitic lifestyle within the genus.

## Acknowledgments

We would like to thank the Consejo Nacional de Investigaciones Científicas y Técnicas (CONICET) for funding the Postdoctoral scholarship, the Instituto Multidisciplinario de Biología Vegetal (IMBIV) for providing the infrastructure to carry out this work and ANPCyT-FONCyT, MINCyT Córdoba, and SECyT-UNC for financial support. We also thank Dr. Mihai Costea from the Department of Biology at the University of Wilfrid Laurier, Waterloo, Canada, for his assistance with species identification.

## References

Andrada AR, Páez VDLA, Toranzo MI, Ruíz De Bigliardo GE. 2018. Análisis de la meiosis y viabilidad del polen en morfotipos intraespecíficos de *Cuscuta paradiana* (Cuscutaceae) [Analysis of meiosis and pollen viability in intraspecific morphotypes of *Cuscuta paradiana* (Cuscutaceae)]. Lilloa. 55(1): 3–15. doi: 10.30550/j.M/2018.55.1/1.

Banerjee A, Stefanović S. 2019. Caught in action: fine-scale plastome evolution in the parasitic plants of *Cuscuta* section *Ceratophorae* (Convolvulaceae). Plant Mol Biol. 100(6):621–634. doi: 10.1007/s11103-019-00884-0.

Banerjee A, Stefanović S. 2020. Reconstructing plastome evolution across the phylogenetic backbone of the parasitic plant genus *Cuscuta* (Convolvulaceae). Bot J Linn Soc. 194(4):423–438. doi: 10.1093/botlinnean/boaa056.

Báez M, Vaio M, Dreissig S, Schubert V, Houben A, Pedrosa-Harand A. 2019. Together but different: the subgenomes of the bimodal *Eleutherine* karyotypes are differentially organized. Front Plant Sci. 10: 1170. doi: 10.3389/fpls.2019.01170.

Biscotti MA, Carducci F, Olmo E, Canapa A. 2019. Vertebrate genome size and the impact of transposable elements in genome evolution. In: Pontarotti P, editor. Evolution, origin of life, concepts and methods. Cham: Springer International Publishing; p. 233–251. doi: 10.1007/978-3-030-30363-1_12.

Biscotti MA, Olmo E, Heslop-Harrison JS. 2015. Repetitive DNA in eukaryotic genomes. Chromosome Res. 23:415–420. doi: 10.1007/s10577-015-9499-z.

Bourque G, Burns KH, Gehring M, Gorbunova V, Seluanov A, Hammell M, Imbeault M, et al. 2018. Ten things you should know about transposable elements. Genome Biol. 19(1):199. doi: 10.1186/s13059-018-1577-z.

Braukmann T, Kuzmina M, Stefanović S. 2013. Plastid genome evolution across the genus *Cuscuta* (Convolvulaceae): two clades within subgenus *Grammica* exhibit extensive gene loss. J Exp Bot. 64(4):977–989. doi: 10.1093/jxb/ers391.

Cabral JS, Felix LP, Guerra M. 2006. Heterochromatin diversity and its co-localization with 5S and 45S rDNA sites in chromosomes of four *Maxillaria* species (Orchidaceae). Genet Mol Biol. 29:659–664. doi: 10.1590/S1415-47572006000400015.

Costea M, García MA, Stefanović S. 2015. A phylogenetically based infrageneric classification of the parasitic plant genus *Cuscuta* (Dodders, Convolvulaceae). Syst Bot. 40(1):269–285. doi: 10.1600/036364415X686567.

Doležel J. 1991. Flow cytometric analysis of nuclear DNA content in higher plants. Phytochem Anal. 2(4):143–154. doi: 10.1002/pca.2800020402.

García MA, Castroviejo S. 2002. Estudios citotaxonómicos en las especies ibéricas del género *Cuscuta* (Convolvulaceae) [Cytotaxonomic studies on the Iberian species of the genus *Cuscuta* (Convolvulaceae)]. An Jard Bot Madrid. 60(1):33–44.

García MA, Costea M, Kuzmina M, Stefanović S. 2014. Phylogeny, character evolution, and biogeography of *Cuscuta* (Dodders; Convolvulaceae) inferred from coding plastid and nuclear sequences. Am J Bot. 101(4):670–690. doi: 10.3732/ajb.1300449.

García MA, Stefanović S, Weiner C, Olszewski M, Costea M. 2018. Cladogenesis and reticulation in *Cuscuta* sect. *Denticulatae* (Convolvulaceae). Org Divers Evol. 18(4):383–398. doi: 10.1007/s13127-018-0378-5.

Goulet BE, Roda F, Hopkins R. 2017. Hybridization in plants: old ideas, new techniques. Plant Physiol. 173(1): 65–78. doi: 10.1104/pp.16.01340.

Gruner A, Hoverter N, Smith T, Knight CA. 2010. Genome size is a strong predictor of root meristem growth rate. J Bot. 2010:1–4. doi: 10.1155/2010/390414.

Guerra M, García MA. 2004. Heterochromatin and rDNA sites distribution in the holocentric chromosomes of *Cuscuta approximata* Bab. (Convolvulaceae). Genome. 47(1):134–140. doi: 10.1139/g03-098.

Heslop-Harrison JS, Schwarzacher T. 2011. Organisation of the plant genome in chromosomes. Plant J. 66(1):18–33. doi: 10.1111/j.1365-313X.2011.04544.x.

Ibiapino A, Báez M, García MA, Costea M, Stefanović S, Pedrosa-Harand A. 2022b. Karyotype asymmetry in *Cuscuta* L. subgenus *Pachystigma* reflects its repeat DNA composition. Chromosome Res. 30(1):91–107. doi: 10.1007/s10577-021-09683-0.

Ibiapino A, García MA, Amorim B, Baez M, Costea M, Stefanović S, Pedrosa-Harand A. 2022a. The evolution of cytogenetic traits in *Cuscuta* (Convolvulaceae), the genus with the most diverse chromosomes in Angiosperms. Front Plant Sci. 13: 842260. doi: 10.3389/fpls.2022.842260.

Ibiapino A, MÁ García, Costea M, Stefanović S, Guerra M. 2020. Intense Proliferation of rDNA Sites and Heterochromatic Bands in Two Distantly Related *Cuscuta* Species (Convolvulaceae) with Very Large Genomes and Symmetric Karyotypes. Genet Mol Biol. 43(3): e20190068.doi: 10.1590/1678-4685-GMB-2019-0068.

Ibiapino A, Urdampilleta J, García MA, Pedrosa-Harand A, Stefanović S, Costea M. 2025. Cytogenetic comparison of *Cuscuta psorothamnensis* and *C. veatchii* (Convolvulaceae), two species originated from recurrent hybridization between the same diploid parents. Plant Syst Evol. 311(2):8. doi: 10.1007/s00606-025-01937-2.

Jhu MY, Sinha NR. 2022. *Cuscuta* Species: Model Organisms for Haustorium Development in Stem Holoparasitic Plants. Front Plant Sci. 13: 1086384. doi: 10.3389/fpls.2022.1086384.

Kaiser B, Vogg G, Fürst UB, Albert M. 2015. Parasitic plants of the genus *Cuscuta* and their interaction with susceptible and resistant host plants. Front Plant Sci. 6:45. doi: 10.3389/fpls.2015.00045.

Kao Y. 2001. Differential accumulation of heterochromatin as a cause for karyotype variation in *Phalaenopsis* orchids. Ann Bot. 87(3):387–395. doi: 10.1006/anbo.2000.1348.

López-Flores I, Garrido-Ramos MA. 2012. The repetitive DNA content of eukaryotic genomes. In: Garrido-Ramos MA, editor. Genome dynamics. Vol. 7. Basel: Karger; p. 1–28. doi: 10.1159/000337118.

Loureiro J, Rodriguez E, Doležel J, Santos C. 2007. Two new nuclear isolation buffers for plant DNA flow cytometry: a test with 37 species. Ann Bot. 100(4):875–888. doi: 10.1093/aob/mcm152.

García MA, Costea M, Guerra M, García-Ruiz I, Stefanović S. 2019. IAPT chromosome data 31. Taxon. 68(6):1374–1380. doi: 10.1002/tax.12176.

McKain MR, Wickett N, Zhang Y, Ayyampalayam S, McCombie WR, Chase MW, Pires JC, De Pamphilis CW, Leebens-Mack J. 2012. Phylogenomic analysis of transcriptome data elucidates co-occurrence of a paleopolyploid event and the origin of bimodal karyotypes in *Agavoideae* (Asparagaceae). Am J Bot. 99(2):397–406. doi: 10.3732/ajb.1100537.

McNeal JR, Kuehl JV, Boore JL, De Pamphilis CW. 2007. Complete Plastid Genome Sequences Suggest Strong Selection for Retention of Photosynthetic Genes in the Parasitic Plant Genus *Cuscuta*. BMC Plant Biol. 7(1): 57. doi: 10.1186/1471-2229-7-57.

Neumann P, Oliveira L, Čížková J, Jang T, Klemme S, Novák P, Stelmach K, Koblížková A, Doležel J, Macas J. 2021. Impact of Parasitic Lifestyle and Different Types of Centromere Organization on Chromosome and Genome Evolution in the Plant Genus *Cuscuta*. New Phytol. 229(4):2365–2377. doi: 10.1111/nph.17003.

Paéz MÁ, Andrada AR, Lozzia ME, Toranzo MI. 2011. Estudios meióticos en cuatro especies del género *Cuscuta* del subgénero *Grammica* (Cuscutaceae) [Meiotic studies in four species of the genus *Cuscuta* of the subgenus *Grammica* (Cuscutaceae)]. Lilloa. 48(1):83–90.

Pan, H., L. Zagorchev, L. Chen, Y. Tao, C. Cai, M. Jiang, Z. Sun, and J. Li. 2023. Complete Chloroplast Genomes of Five *Cuscuta* Species and Their Evolutionary Significance in the *Cuscuta* Genus. BMC Genom. 24(1):310. doi: 10.1186/s12864-023-09427-w.

Pedrosa A, Sandal N, Stougaard J, Schweizer D, Bachmair A. 2002. Chromosomal Map of the Model Legume *Lotus Japonicus*. Genetics 161(4):1661–1672. doi: 10.1093/genetics/161.4.1661.

Piednoël M, Aberer AJ, Schneeweiss GM, Macas J, Novak P, Gundlach H, Temsch EM, Renner MM. 2012. Next-Generation Sequencing Reveals the Impact of Repetitive DNA Across Phylogenetically Closely Related Genomes of Orobanchaceae. Mol Biol Evol. 29(11):3601–3611. 10.1093/molbev/mss168.

Roa F, Guerra M. 2012. Distribution of 45S rDNA Sites in Chromosomes of Plants: Structural and Evolutionary Implications. BMC Evol Biol. 12(1):225. 10.1186/1471-2148-12-225.

Roa F, Guerra M. 2015. Non-Random Distribution of 5S rDNA Sites and Its Association with 45S rDNA in Plant Chromosomes. Cytogenet Genome Res 146(3):243–249. doi: 10.1159/000440930.

Van-Lume B, Esposito T, Diniz-Filho JAF, Gagnon E, Lewis GP, Souza G. 2017. Heterochromatic and Cytomolecular Diversification in the Caesalpinia Group (Leguminosae): Relationships between Phylogenetic and Cytogeographical Data. PPEES. 29:51–63. doi: 10.1016/j.ppees.2017.11.004.

Van-Lume B, Mata-Sucre Y, Báez M, Ribeiro T, Huettel B, Gagnon E, Leitch IJ, Pedrosa-Harand A, Lewis GP, Souza G. 2019. Evolutionary Convergence or Homology? Comparative Cytogenomics of Caesalpinia Group Species (Leguminosae) Reveals Diversification in the Pericentromeric Heterochromatic Composition. Planta. 250(6):2173–2186. 10.1007/s00425-019-03287-z.

Vimala Y, Lavania S, Lavania UC. 2021. Chromosome Change and Karyotype Differentiation–Implications in Speciation and Plant Systematics. The Nucleus. 64(1):33–54. doi: 10.1007/s13237-020-00343-y.

Waminal NE, Pellerin RJ, Kim NS, Jayakodi M, Park JY, Yang TJ, Kim HH. 2018. Rapid and Efficient FISH Using Pre-Labeled Oligomer Probes. Sci Rep. 8(1):8224. doi: 10.1038/s41598-018-26667-z.

